# BugBuster: A novel automatic and reproducible workflow for metagenomic data analysis

**DOI:** 10.1101/2025.02.24.639915

**Authors:** Francisco Fuentes-Santander, Carolina Curiqueo, Rafael Araos, Juan A. Ugalde

**Author notes:** Corresponding author. Juan A. Ugalde.

## Abstract

**Summary:** In metagenomic sequencing, large volumes of data are obtained with all the genetic information present in a sample, allowing valuable data to be obtained about microbial communities. The software and processes necessary to obtain quality results have become increasingly complex and sophisticated, limiting the accessibility of biologists who try to use them. To facilitate the analysis of this data, a modular and reproducible workflow was developed using the Nextflow workflow orchestrator named BugBuster. The pipeline is easy to implement because all its dependencies are provided within containers, it is reproducible, modular and portable. BugBuster consists of different processes that allow data analysis at the level of reads, contigs and MAGs, also including modules for resistome characterization and taxonomic profiling.

**Availability and implementation:** BugBuster was written in Nextflow DSL2 Syntaxis. The program applications, user manual, exemplary data and code are freely available at https://github.com/gene2dis/BugBuster.

## 1 Introduction

Metagenomics has established itself as a fundamental tool in diverse areas of microbiology, including pathogen identification, antimicrobial resistance monitoring, and industrial process optimization (Bashir et al., 2014).

Processing a metagenomic sample involves multiple steps, such as removing low-quality sequences and contaminants, taxonomic profiling, genome assembly, gene annotation, and binning. While various tools exist to perform each step, their outputs and standards are often incompatible or not directly comparable, adding complexity to the analysis of metagenomic datasets (Tamames & Puente-Sánchez, 2018). The large volumes of data generated require not only advanced computational tools but also expertise in high-performance infrastructure to run large-scale analyses efficiently. Numerous pipelines and workflows have been developed to process microbial data following specific protocols for different analytical stages (Navgire et al., 2022). Nevertheless, experienced users frequently assemble custom hybrid workflows by selecting analysis that best fit their requirements, adding complexity and challenges to reproducibility.

Despite their flexibility, these hybrid approaches pose significant challenges owing to the lack of standardized interoperability between tools, which requires manual adjustments to integrate inputs and outputs. Additionally, the various and evolving dependencies associated with these workflows make them difficult to implement, even for experienced users, and are nearly inaccessible to beginners, particularly in unfamiliar computational environments (Kesh & Raghupathi, 2004). Updating these workflows to incorporate new methodologies often requires rewriting scripts, further complicating maintenance and usability.

In addition to these concerns regarding interoperability and usability, reproducibility has become a key concern in computational biology. Publishing data and scripts on platforms such as GitHub or GitLab, are becoming standard practice and even a requirement in some cases. However, making these resources available does not guarantee reproducibility because platform-specific dependencies can hinder deployment across different environments (Baykal et al., 2024). In addition, differing versions of libraries and tools on HPC systems can further complicate reproducibility.

A potential solution to these challenges is the use of container applications, such as Docker, which allow users to run applications in isolated environments by packaging all the necessary components, including files, libraries, and operating systems (Docker, 2020). When combined with a workflow management system, such as Nextflow, these containers enable the creation of automated, scalable, reproducible, efficient, and portable workflows (Di Tommaso et al., 2017).

To address these challenges, we present BugBuster, a fully automated workflow for metagenomic data processing that covers all stages of analysis, from initial quality control to resistome detection and characterization. BugBuster was developed in Nextflow using the DSL2 syntax, providing a modular and flexible structure. Each process was encapsulated in Docker containers to ensure easy installation and high reproducibility. Additionally, it includes specialized modules for identifying antibiotic resistance genes that can be directly associated with specific taxa.

## 2 BugBuster pipeline

BugBuster is written in Nextflow using the DSL2 syntax, which allows for a modularized pipeline structure. It consists of 61 modules that facilitate customization during execution. These 61 modules can be grouped into six steps that address metagenome processing: i) Reads processing, ii) Taxonomic profiling reads, iii) Prediction of of antibiotic resistance genes (ARGs) and resistance-causing gene variants (ARGVs) in reads, iv) Assembly; v) Binning; vi) Taxonomy prediction and resistance genes in contigs. Some of the customization options currently available are:

1. Assembly mode: Assembly, all metagenomes were processed individually; co-assembly and metagenomes were grouped, and a single assembly was performed.
2. Software used for taxonomic profiling: Kraken2 (Wood et al., 2019) and Sourmash (Titus Brown & Irber, 2016).
3. Prediction of resistance genes and gene variants that cause antibiotic resistance in reads.
4. Functional annotation, taxonomic prediction, and resistance genes prediction from contigs.
5. Metagenomic binning

### 2.1 Description of BugBuster modules

#### 2.1.1 Read processing

Read quality filtering is performed using FastP v. 0.23.2 (Chen et al., 2018) with the following parameters --unqualified_percent_limit=10, --cut_front, --cut_front_window_size=4, --cut_front_mean_quality=20, --cut_right, --cut_right_window_size=4, --cut_right_mean_quality=20, --detect_adapter_for_pe, --n_base_limit=5 and --trim_poly_g these are the parameters set by default to BugBuster. Subsequently, the samples containing the minimum number of reads specified by the user are filtered. Additionally, contaminating reads are discarded by mapping against the human reference genome T2T-CHM13v2.0 (Accession number: GCA_000001405.1) and the PhiX genome (Accession number: NC_001422) with Bowtie2 v. 2.5.3 (Langmead & Salzberg, 2012) using the parameters -N=1, -L=20, -score-min=‘G,15,6’, -R=2, -i ‘S,1,0.75’. Data from the filtered reads during all steps are collected using a Bash script, and a report and plots are generated using an R script.

#### Resistance gene prediction in reads

Prediction of ARGs and ARGVs at the read level is performed with KARGA v. 1.02 (Prosperi & Marini, 2021) and KARGVA v, 1.0 (Marini et al., 2023) both using a kmer length of 17. The predicted ARG genes are filtered by 90%>= gene coverage, and ARGV genes are filtered by 80%>= gene coverage and with at least 2 KmerSNPHits. Predicted genes are normalized by estimating the number of cells with ARGs-OAP v.

3.2.4 (Yin et al., 2023) using the default options. All the generated results are unified in a report using an R script.

#### 2.1.3 Taxonomic profiling in reads

Kraken2 workflow: Taxonomic prediction and abundance estimation at the read level is performed using Kraken2 v. 2.1.3 in conjunction with Bracken v. 2.9 (Lu et al., 2017). The results generated by Bracken are unified using Kraken-Biom v. 1.2.0 (Dabdoub, 2016). Subsequently, the generated file in the biom format is transformed into a Phyloseq object (McMurdie & Holmes, 2013).

##### Sourmash workflow

An alternative to Kraken2, is Sourmash, which uses MinHash sketches to represent the taxonomic signatures of large sequence sets, allowing for more efficient storage, reducing RAM and CPU usage, and allowing more extensive reference databases. Taxonomic prediction and abundance estimation are performed using Sourmash v. 4.8.11. Estimation of the k-mer content in the reads is performed using the Sourmash *sketch dna* function with different parameters depending on the requested taxonomic resolution: Genus: -p k=21,scaled=1000,abund; Species: -p k=31,scaled=1000,abund; Strain: -p k=51,scaled=1000,abund. The minimum metagenome coverage estimation is then performed using the Sourmash *gather function* with its default parameters, changing the k-mer length according to the requested taxonomic resolution (Genus: k21; Species: k31; Strain: k51). Subsequently, the Sourmash *tax annotation* function is used to obtain the taxonomy assigned to the reads, and the generated file is used to create a Phyloseq object with an R script.

Finally, data on the proportion of taxonomically classified reads is collected for both workflows using a Bash script. These results are used to generate relative abundance plots using the microViz package v. 0.12.3 (Barnett et al., 2021) and bar plots with the proportion of classified reads using *ggplot2* v. 3.5.0

#### 2.1.4 Assembly

Read assembly is performed using MegaHit v. 1.2.9 (Li et al., 2015), in two different modes: per sample, where each metagenome is processed individually; and co-assembly, where all the samples are grouped, and a single assembly is performed. Contigs smaller than 1,000 bp are filtered using BBmap version 39.06 (Bushnell, 2014), and a report is generated with assembly metrics.

#### 2.1.5 Taxonomic annotation and antibiotic resistance genes identification in contigs

Taxonomic annotation of contigs is first performed with a search using Blastn v. 2.15.0 (Altschul et al., 1990) against the NT database. Then, the search result is used to assign taxonomy with BlobTools v.1.1.1 (Laetsch & Blaxter, 2017) using the default parameters. Identification of antibiotic resistance genes at the contig level is performed with DeepARG v. 1.0.4 (Arango-Argoty et al., 2018) using the default parameters. The results generated with both software are unified in a CSV file and visualized with an R script.

#### 2.1.6 ORF Prediction and Functional Annotation in Contigs

ORF prediction in contigs is performed using Prodigal v. 2.6.3 (Hyatt et al., 2010) with the metagenomics option. Functional annotation at the contig level is performed using MetaCerberus v. 1.2.1 (Figueroa et al., 2024), using, by default, the KOFam_all, COG, VOG, PHROG, and CAZy HMMs available in its database.

#### 2.1.7 Binning and bin refinement

Metagenome binning is performed using Metabat2 v. 2.15 (Kang et al., 2019), Semibin2 v. 2.1.0 (Pan et al., 2023) using the human-intestine trained model by default (modifiable by the user) and Comebin v. 1.0.4 (Wang et al., 2024) with 3 attempts, the first with the default options, followed by reduction of the embedding size in case there is a failure on the process, using the following parameters: -b 896, -e 1792, -c 1792 for the second attempt, and -b 512, -e 1024, -c 1024 for a final attempt. Subsequently, the MAGs generated are refined with MetaWrap v.1.2 (Uritskiy et al., 2018), using default thresholds of a minimum completeness of 50% and a maximum contamination of 10%.

#### 2.1.8 Quality estimation and taxonomic prediction of bins

The quality prediction of unrefined and refined bins is performed with CheckM2 v. 1.0.1 (Chklovski et al., 2024) using the default parameters. The refined bins are taxonomically classified using GTDB-TK v. 2.4.0 (Chaumeil et al., 2022) against the GTDB database release 220. Finally, the results are unified in a unique CSV file, which is used to generate quality and taxonomy graphs using an R script. In the co-ensemble mode, the coverage in each sample is also calculated separately for each refined bin using Bedtools v. 2.31.1 (Quinlan & Hall, 2010) and Samtools v. 1.17 (Danecek et al., 2021), and the coverage data are unified with the taxonomy and quality data in a CSV file.

## 3 Tests dataset

To illustrate the results of the BugBuster pipeline, we used 9 simulated samples of metagenomic data from the human gastrointestinal tract from CAMI (Fritz et al., 2019). Using this data, we tested the pipeline with these options: --assembly_mode “assembly” --taxonomic_profiler “sourmash” --read_arg_prediction --contig_tax_and_arg --include_binning. All the data was processed on an 80-CPU Intel Xeon E7-4820 v4 server with 2TB RAM.

## 4 Results

### 4.1 Reads preprocessing

The processing of simulated gut microbiota reads shows a progressive decrease in the total number of reads throughout the filtering steps. A total of 1.62% of reads were removed based on quality criteria, and no human contamination was detected (Figure S1).

### 4.2 Taxonomic profiling and abundance estimation

The results show a comparison between Sourmash and Kraken using the GTDB release 207 database and the simulated data. On average, Kraken classified approximately 96.5% of the reads, whereas Sourmash classified approximately 62.5% (Figure S2). We kept the proportions of taxa in the two workflows with respect to the CAMI data set at the phylum level, observing differences only in the taxonomic names assigned by the respective databases due to different naming conventions between NCBI and GTDB (Figure S3). In particular, variations in the naming of taxonomic groups within the phylum Firmicutes are mainly due to the number of genomes included in the GTDB database and their detailed taxonomic classification at the strain level (Parks et al., 2022). As a result, the GTDB database assigns additional identifiers, such as letter codes, after the phylum name (e.g., Firmicutes_B) to distinguish the genomes of unique species.

### 4.3 Resistance gene prediction in reads

Processed reads can also be utilized to predict antibiotic resistance genes (ARGs) and their variants (ARGVs). For ARG prediction, we used the Megares v3.0 database (Bonin et al., 2023), identifying 824 genes in total (Table S1). For ARGV prediction, we employed the KARGVA v5 database (Marini et al., 2023), which led to the identification of 358 predicted genes in total (Table S2).

### 4.4 Assembly, taxonomic prediction, and resistance gene prediction in contigs

Each sample was assembled individually, and contigs larger than 1 KB were kept for further analysis. Results show the taxonomy of the contigs present in all samples using blobplots, we obtained an average N50 of 2.175 KB and taxonomically classified 99.88% of the generated contigs throughout the entire set of samples (Figure 2, top and bottom panel). The proportion of phyla in the contigs obtained by BugBuster was similar to that provided by the simulated CAMI data, with Bacillota being the most abundant phylum (Table S3). All filtered contigs were used to search for resistance genes using DeepARG identifying 1706 predicted genes throughout the entire set of samples (Figure S6).

**Figure 1.**
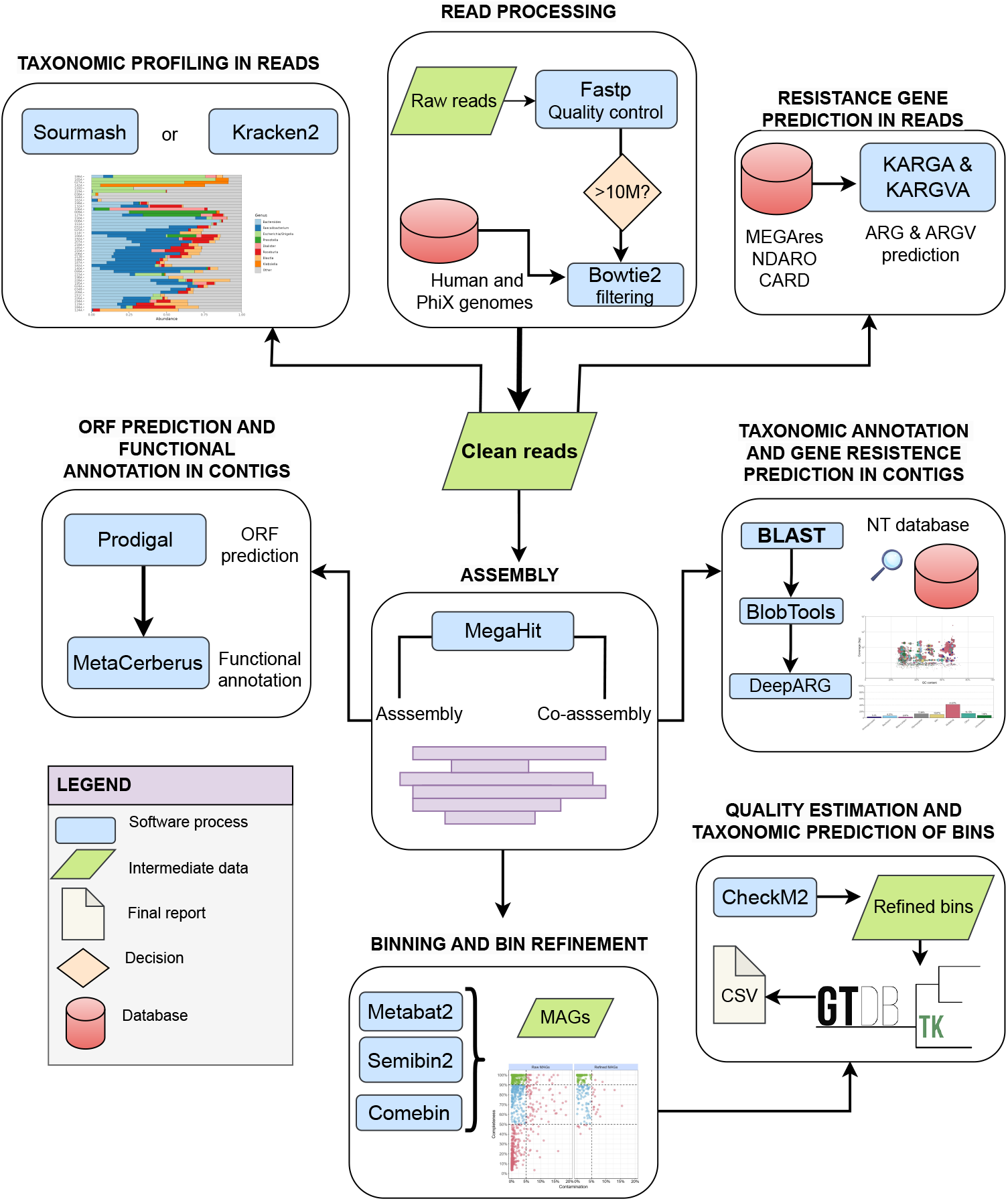
Workflow of the modules and customizable decisions during execution. The execution performs: 1) Quality filtering of reads; 2) Filtering of samples containing a minimum number of reads specified by the user; 3) Filtering of contaminant reads; 4) Prediction of antibiotic resistance genes and gene variants causing resistance at the read level; 5) Normalization of predicted genes by estimating the number of cells; 6) Taxonomic prediction and abundance estimation at the read level; 7) Reports for tracking reads and taxonomic reports; 8) Read assembly; 9) Taxonomic annotation of contigs; 10) Functional annotation of contigs; 11) ORF prediction in contigs; 12) Prediction of resistance genes at the contig level; 13) Contig report and two-dimensional scatter plots; 14) Contig filtering; 15) Metagenomic binning; 16) Refinement of assembled genomes in metagenomes (MAGs); 17) Prediction of MAG quality; 18) Taxonomic prediction of MAGs; 19) MAG results report.

**Figure 2.**
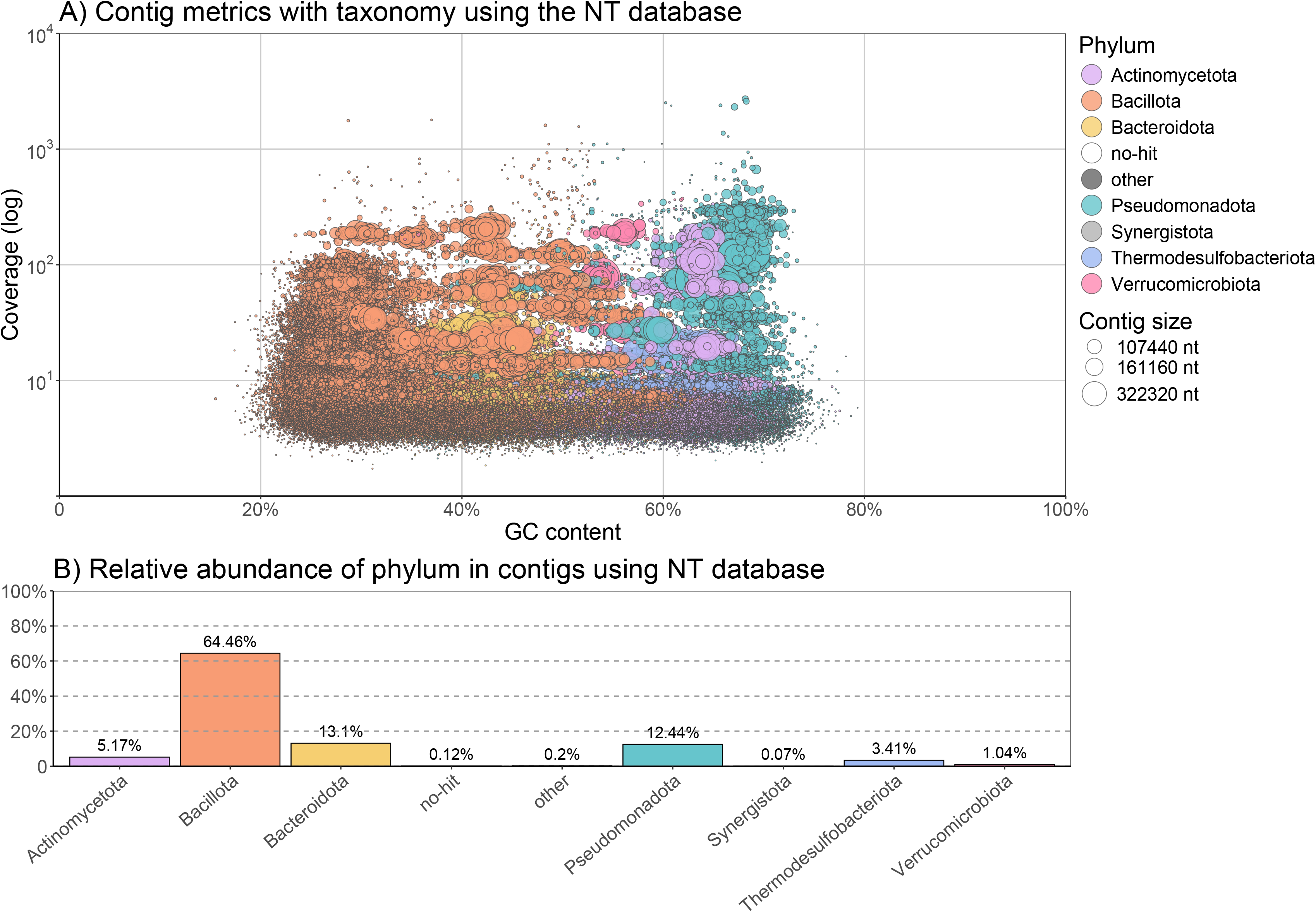
Summary of the metrics obtained during the taxonomic classification of the contigs across all sample sets. A) Blobplot with contig metrics and taxonomy assigned using Blastn, Blobtools and the NT database. B) Relative abundance of the taxonomy at phylum level in the contigs, classified with Blastn, BlobTools, and the NT database.

### 4.5 Binning, quality estimation, and taxonomic prediction in bins

For binning, we obtained 112, 72, and 95 high-quality bins; 61, 66, and 73 medium-quality bins; and 221, 135, and 169 low-quality bins from Comebin, MetaBAT2, and SemiBin2, respectively. This set of bins was used to generate 115 high-quality, 73 medium-quality, and 27 low-quality bins with MetaWRAP (Figure S4). In total, we reconstructed 215 metagenome-assembled genomes (MAGs) of 266 genomes provided in the CAMI dataset. All these 215 MAGs were successfully classified taxonomically at species level (Figure S5).

## 5 Conclusions

BugBuster provides an easy-to-implement and highly reproducible workflow covering pre-processing, taxonomic classification, antibiotic resistance gene prediction, assembly, and MAG refinement. It includes documentation for users (https://github.com/gene2dis/BugBuster) and generates visual outputs for a better interpretation of the results. BugBuster offers modular flexibility, allowing users to select specific modules and optimize configurations for their research needs. The pipeline will be continuously updated to integrate the latest analysis methods. Its DSL2-based modularity enables the efficient incorporation of new tools, including future enhancements for mobile genetic element detection, resistance gene clustering, and optimized execution on limited computational resources.

## Supplementary data

**Figure S1**. Tracking of the reads at each filtering step. Bowtie PhiX, and Bowtie Human represent the number of reads that passed the filtering for PhiX phage and human reads, respectively.

**Figure S2**. Proportion of reads taxonomically classified with both read taxonomic classification workflows. A) Reads classified with kraken 2 with confidence of 0.1 and GTDB release 207. B) Reads classified with sourmash at species-level configuration and GTDB release 207.

**Figure S3**. Relative abundance of the simulated gut microbial communities. A) Relative abundance for the simulated communities, provided in CAMI challenge. B) Relative abundance with BugBuster, using Kraken 2 with confidence of 0.1 and GTDB release 207. C) Relative abundance with BugBuster, using Sourmash at species level configuration and GTDB release 207.

**Figure S4**. Quality evaluation of the generated MAGs. Raw MAGs refers to the MAGs generated with Comebin, Semibin2, and Metabat2 across all samples. Refined MAGs represent the total number of refined MAGs reconstructed across all samples.

**Figure S5**. Summary of the phylum-level taxonomy of the MAGs generated for the entire sample set.

**Figure S6**. Summary of the contigs metrics mixed with ARG predictions across all sample sets. A) Blobplot with contig metrics and ARGs predicted using Deeparg. B) Relative abundance of ARGs found in contigs using Deeparg.

**Table S1** BugBuster merged results from KARGA and ARGs-OAP.

**Table S2** BugBuster merged results from KARGVA and ARGs-OAP.

**Table S3** Comparison between BugBuster predicted taxonomy of contigs vs real taxonomy provided by CAMI dataset

