## Supplementary figures and images for "BugBuster: A novel automatic and reproducible workflow for metagenomic data analysis"

### Supplemental Figure 1

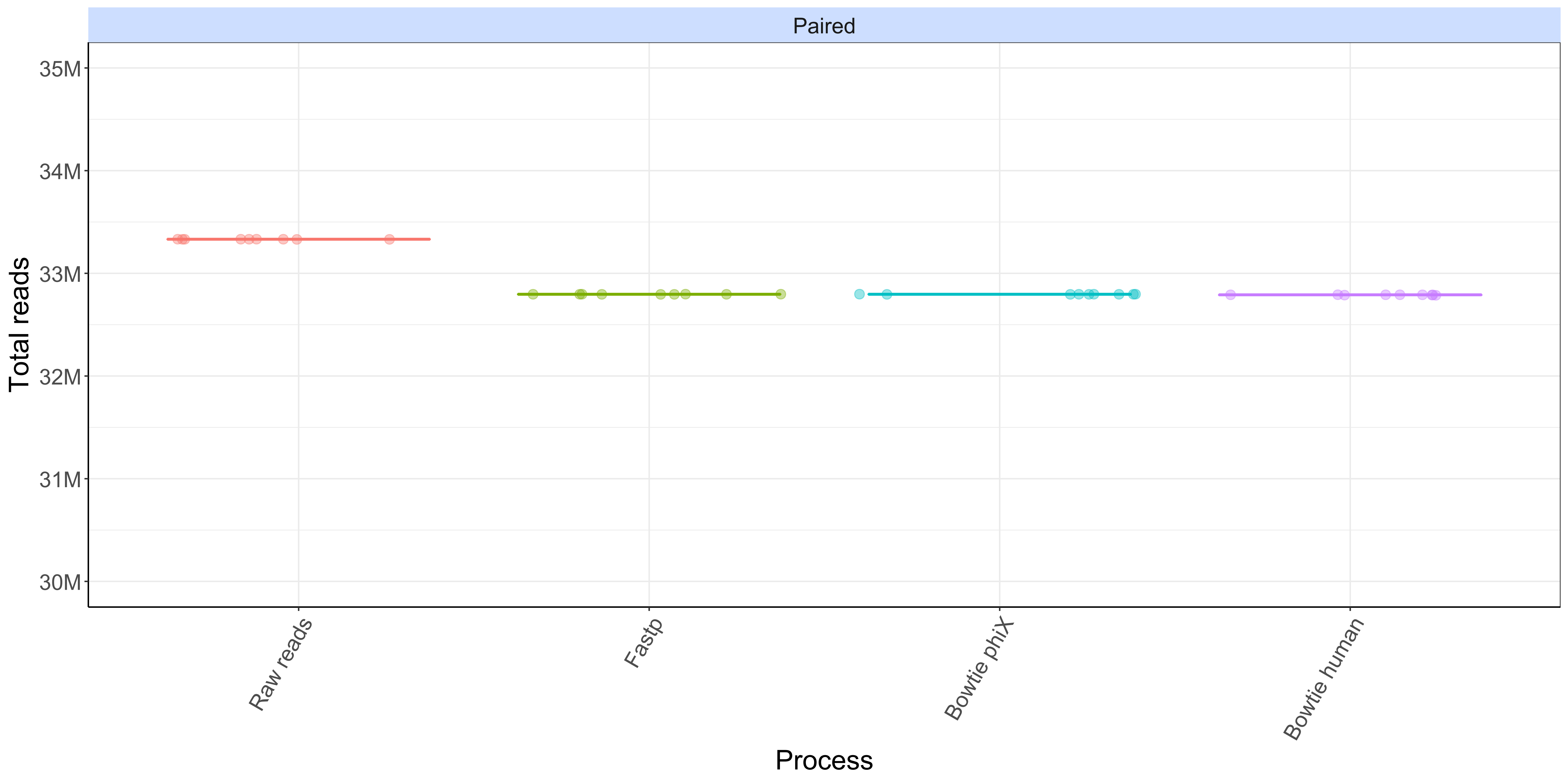

### Supplemental Figure 2

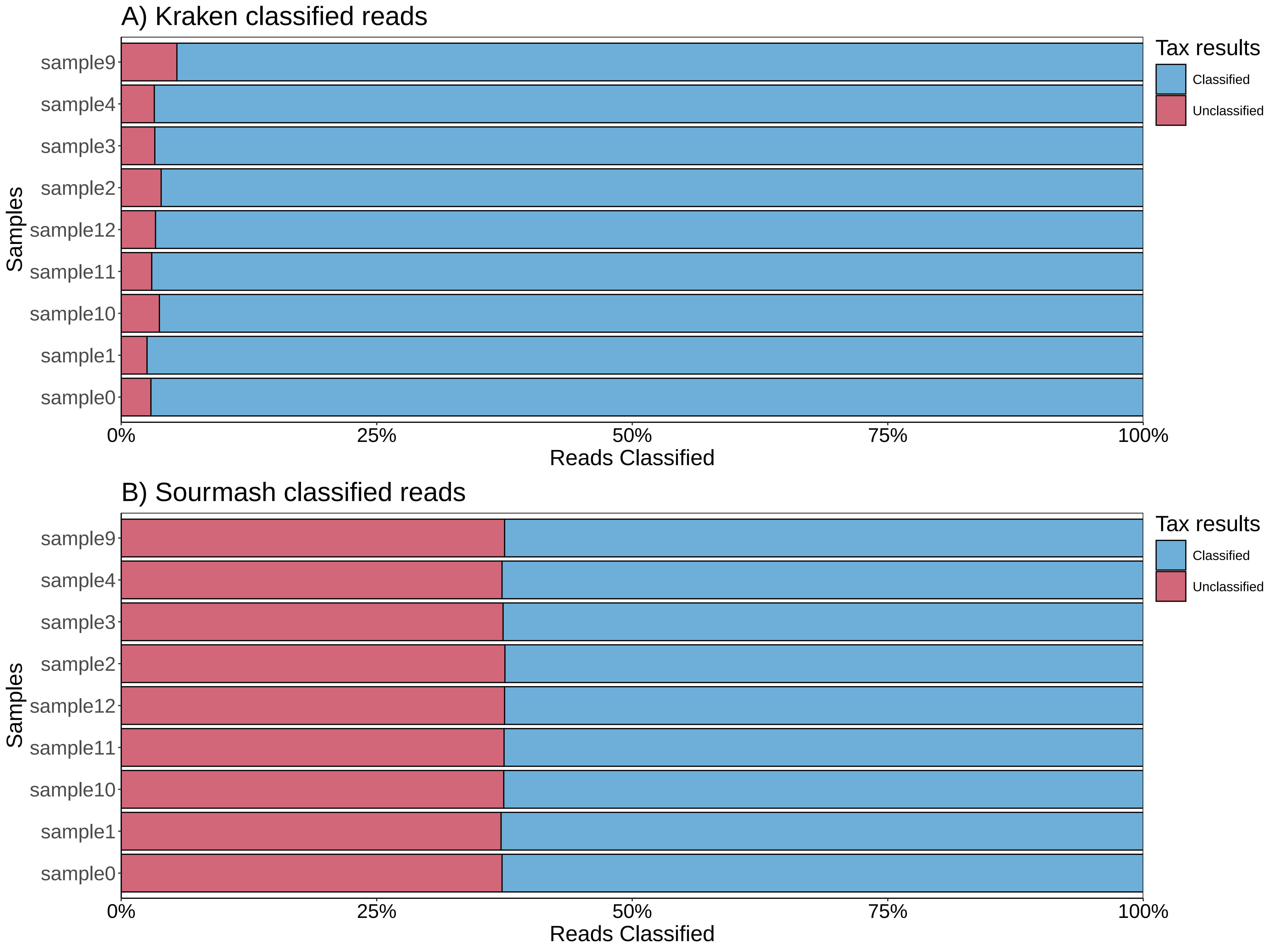

### Supplemental Figure 3

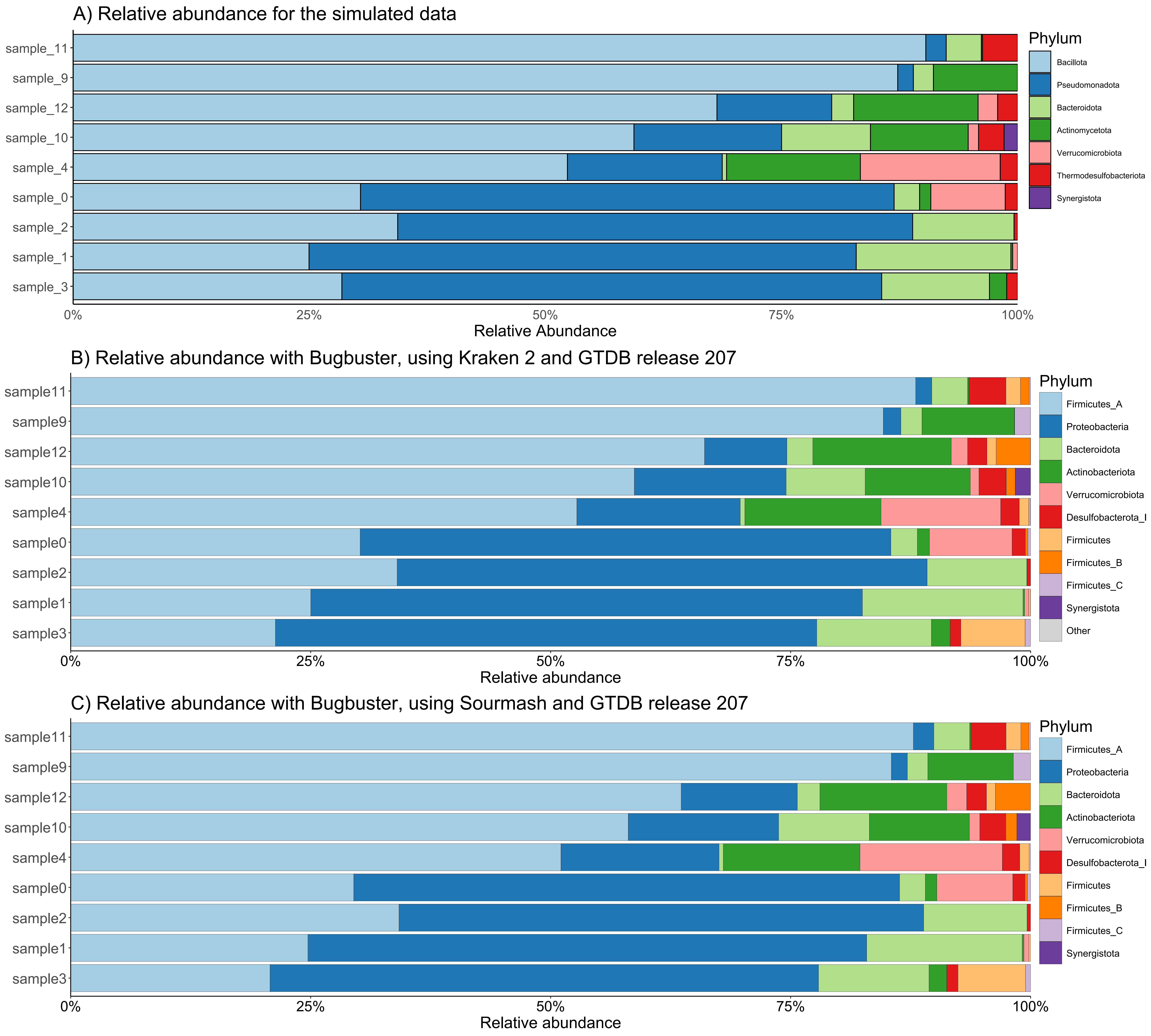

### Supplemental Figure 4

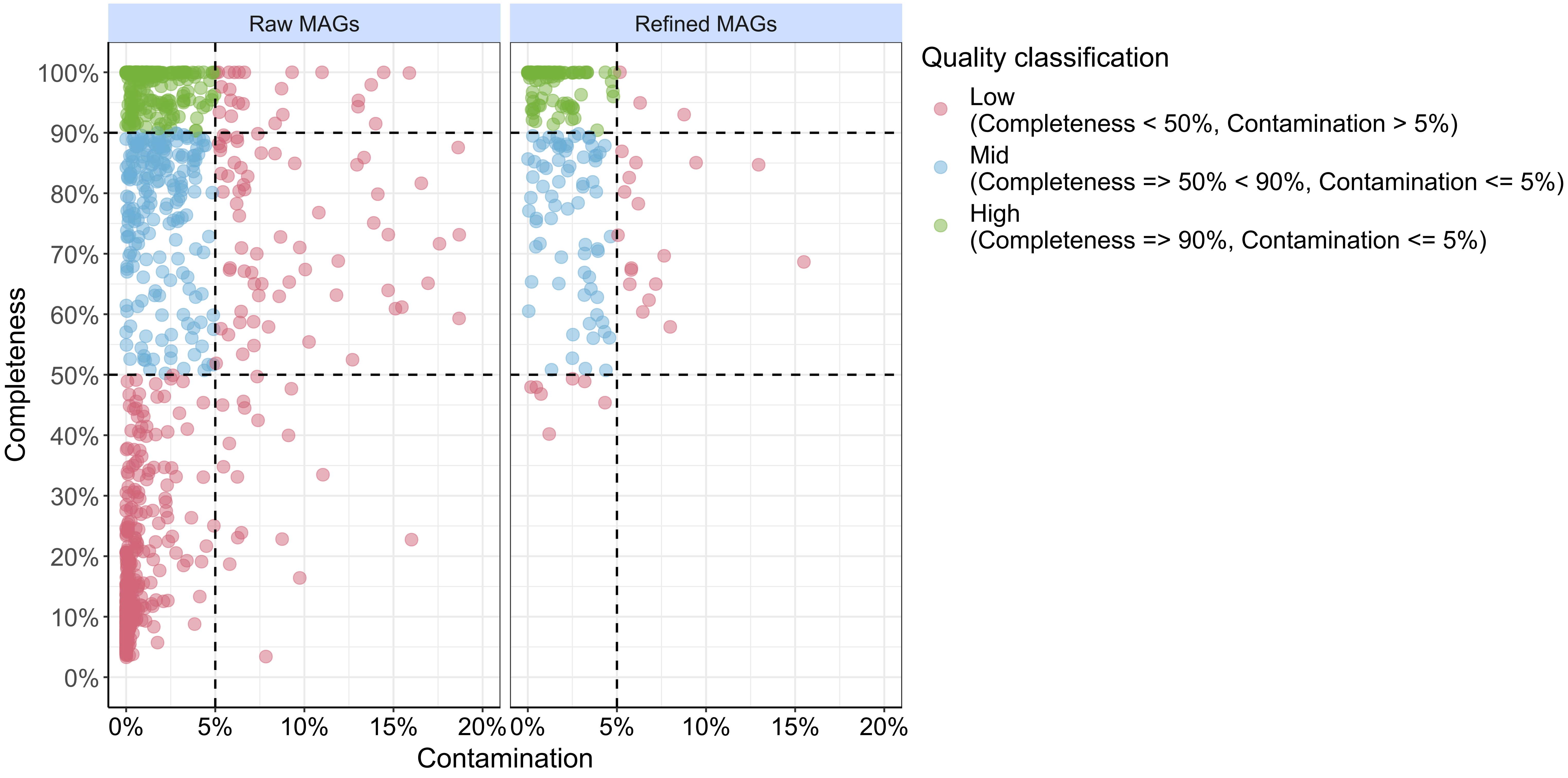

### Supplemental Figure 5

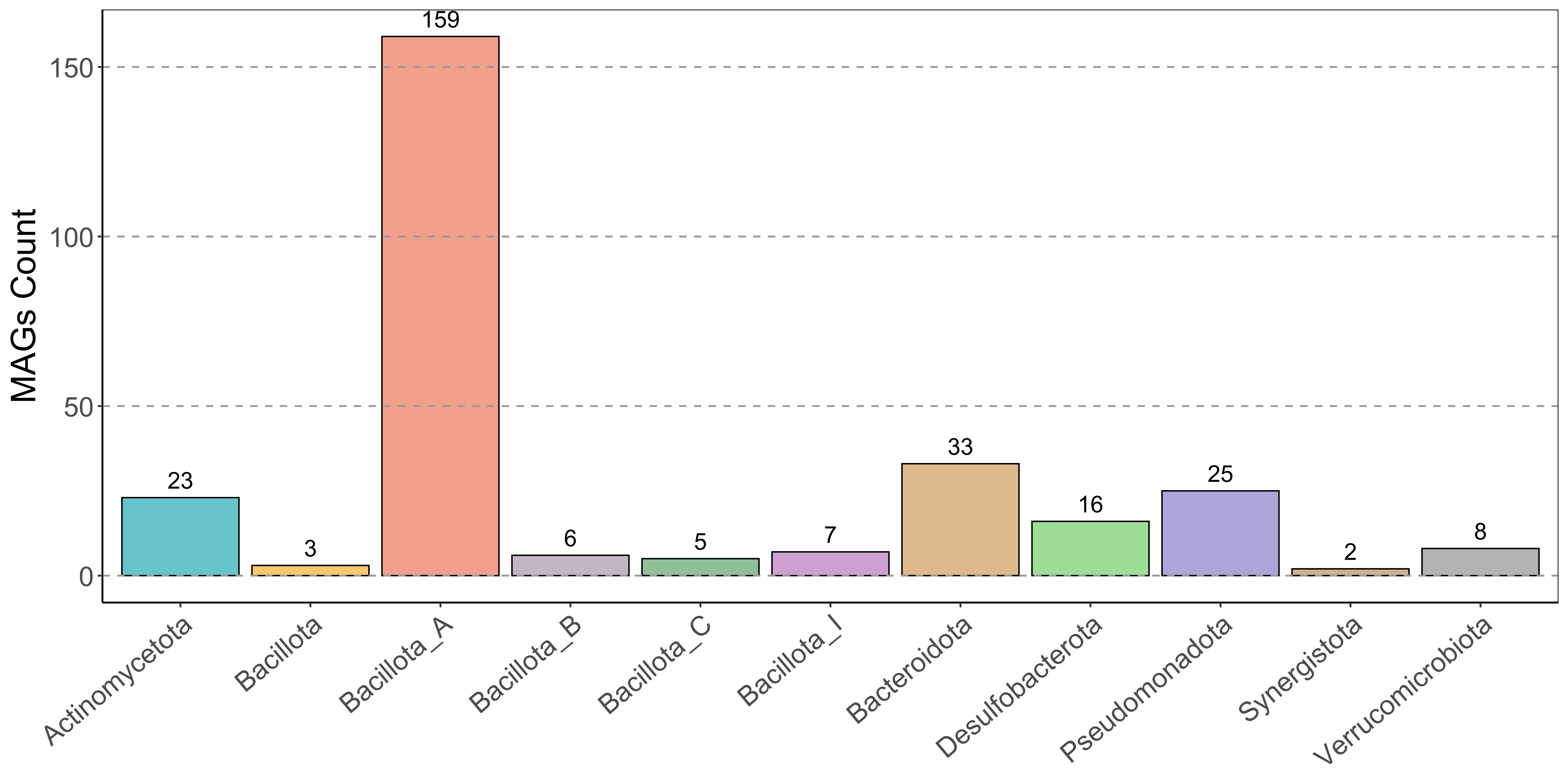

### Supplemental Figure 6

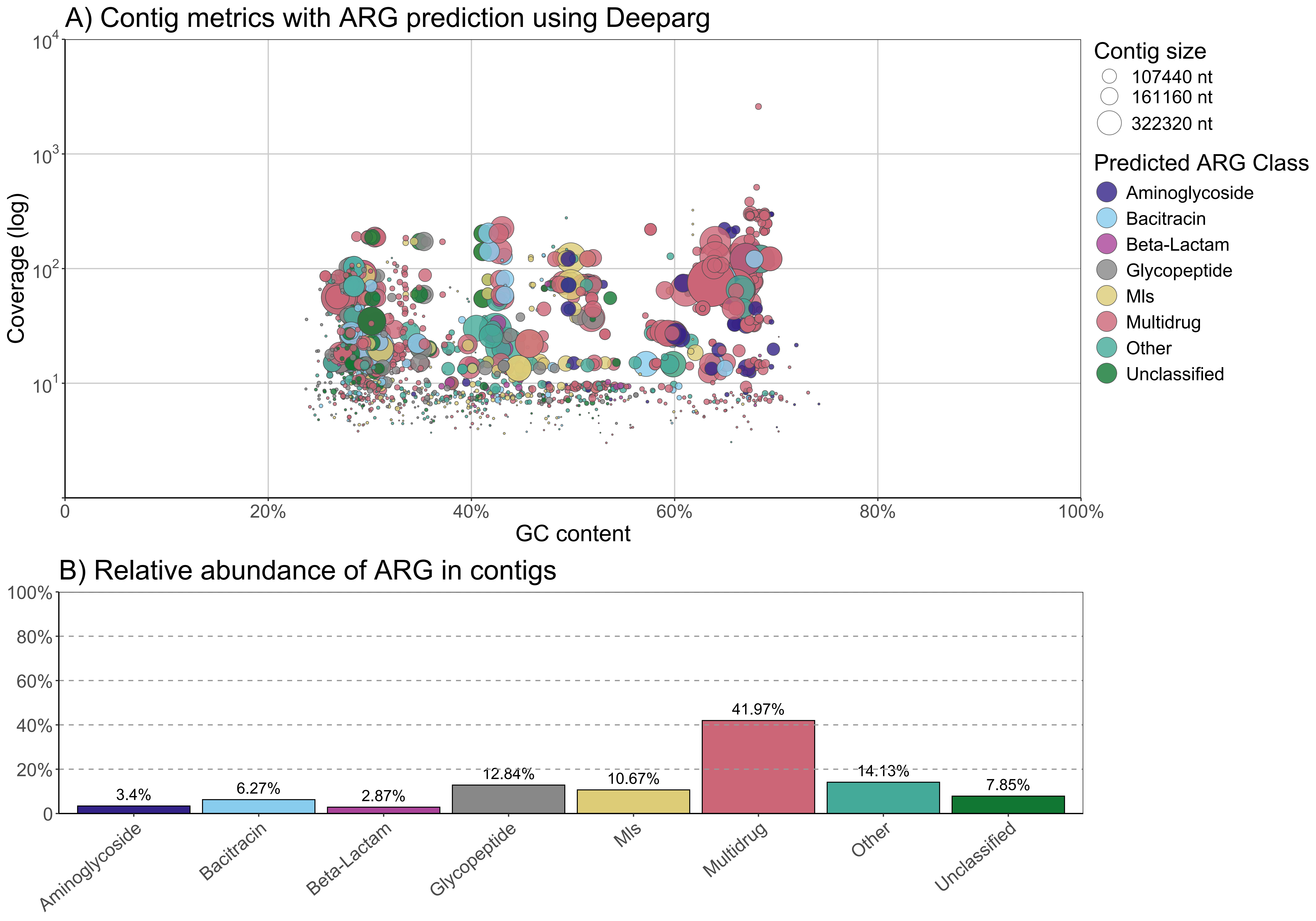
